# Neural reinstatement guides context-dependent emotional memory retrieval

**DOI:** 10.1101/785899

**Authors:** Augustin C. Hennings, Mason McClay, Jarrod A. Lewis-Peacock, Joseph E. Dunsmoor

## Abstract

Reactivating the memory of a context improves retrieval of information learned in that context. But does context reactivation resolve memory competition between related but conflicting emotional experiences? Here, we asked whether spontaneously retrieved episodic context disambiguates competing memories of fear and safety in the healthy brain and in posttraumatic stress disorder. During fMRI, subjects learned that items from an object category were a threat, and then learned that different items from the same category were safe in a unique context. The next day, subjects viewed new threat ambiguous stimuli from the same category and reported their expectation of threat. Multivoxel pattern analysis (MVPA) was used to identify and decode neural patterns unique to the context of safety learning. We show that in healthy adults the degree of neural reinstatement of the safe context predicted activity to threat ambiguous stimuli in the ventromedial prefrontal cortex (vmPFC) and hippocampus, regions important for contextual modulation of emotional memory retrieval. The degree of context reinstatement also disambiguated subjects’ feelings of threat versus safety. In contrast, context reinstatement did not resolve emotional memory retrieval in subjects with PTSD, indicating a contextual processing deficit in this group. Finally, multivoxel pattern similarity between safe and threat ambiguous stimuli overlapped in the vmPFC in healthy adults but not in PTSD. These findings reveal how neural reinstatement helps resolve context-dependent memory retrieval between opposing sources of emotional information. Understanding how the human brain separately encodes and retrieves competing memories of related events has broad implications for the study of emotional memory, associative learning, and the neuropathophysiology of affective disorders.

---

An adaptive memory system should be capable of maintaining conflicting memories of related experiences, and retrieving the appropriate memory given the current circumstances. How neural competition between conflicting memories is resolved remains an important question. Consider for example how you may feel about seafood after eating a dish containing tainted seabass while on a summer vacation. Nonetheless, you may later consume salmon at a local restaurant and find it delightful with no aversive consequences. This second experience countervails previous learning, and creates a new representation that “seafood is safe.” These two opposing learning events may enter into conflict when there is an opportunity to eat a new seafood dish: does this meal pose a threat or is it harmless? One way the brain resolves ambiguity in situations like this is by retrieving past memories of similar situations^1^. Episodic memories, for instance, involve event-specific details embedded in the contextual information present during the time of the original experience^2^. Thinking about a past event can bring its context to mind, and vice versa. Consequently, retrieving contextual details of either the aversive or safe dining experience could bias memory retrieval in favor of either an aversive or a safe association with seafood, thereby inhibiting expression of the alternative association^3^. This same scenario plays out in more extreme emotional situations as well. For example, posttraumatic stress disorder (PTSD) is characterized in part by the inability to inhibit memories of threat in harmless environments^4^. Here, we investigated whether neural reinstatement of a past context resolves context-dependent emotional memory retrieval during time of threat ambiguity in healthy adults and participants with PTSD.

To address this question, we leveraged theoretical insights and experimental approaches from two academic traditions that seldom intersect: episodic memory and Pavlovian conditioning. Episodic memories are easier to recall if the context at retrieval matches that from encoding, sometimes referred to as the encoding-retrieval match or transfer appropriate processing ^5^. Background spatiotemporal details from the time of episodic memory formation can also provide a “mental context,” and reinstatement of the mental context can help guide memory retrieval^6^. Neuroimaging experiments using multivoxel pattern analysis (MVPA) have cleverly incorporated the concept of mental context to decode brain activity related to the retrieval of items that had been encoded in a perceptually unique context^7–9^. Whether a mental context framework can be applied to understand context-dependent emotional memory retrieval is unknown.

Pavlovian conditioning is an ideal model to examine how context resolves memory retrieval between two related, but incompatible emotional experiences. Experimental studies of conditioned fear and clinical accounts of PTSD demonstrate that extinction memories are contextually specific, and fear often returns outside the extinction context in a form of fear relapse known as renewal^1,10^. Extinction, the subsequent learning of safety following fear conditioning, generates a secondary memory that inhibits retrieval and expression of the original threat memory. But extinction also introduces ambiguity to the emotional meaning of a conditioned stimulus^3^, because the stimulus can now signal either threat or safety. The context at the time of memory retrieval helps resolve this ambiguity and determines which memory (threat or safety), and consequent behavior, is most appropriate^1^. Whereas memory of threat transverses environments, memory of the countervailing experience of safety is bound to the context where safety was learned. Computations in and between the ventromedial prefrontal cortex (vmPFC), hippocampus, and amygdala modulates context-dependent extinction memory retrieval^11^. Importantly, while research utilizing Pavlovian conditioning often defines context in terms of the physical environment (i.e., an animal’s cage), conditioning models account for a variety of contextual cues that include spatiotemporal factors and internal bodily or mental states^12^.

Here, we asked whether neural reinstatement of an extinction mental context reveals the context-dependent nature of emotional memory retrieval in humans. We generated a mnemonic signature specific to the encoding of extinction using MVPA tools appropriate for detecting context-dependent memory reactivation. This was combined with a novel Pavlovian conditioning protocol in order to track reinstatement of an extinction context signature during a 24-hour delayed test of fear renewal, in which subjects encountered unique threat ambiguous exemplars that belonged to a semantic category of objects used during both fear conditioning and extinction the previous day. (Fig 1A). We compared results between healthy adults and individuals with PTSD, a disorder characterized by severe dysregulation in contextual processing that might contribute to deficits in the retrieval of extinction memories^4^. We specifically tested whether presentations of unique threat ambiguous stimuli trigger neural reinstatement of the specific mental context in which extinction was learned to facilitate retrieval of extinction over the retrieval of fear. We therefore predicted that healthy adults, but not individuals with PTSD, would show a positive correspondence between the degree of extinction mental context reinstatement and activation within canonical extinction neurocircuitry at a 24-hour test of fear renewal. In addition to probing the contextual nature of emotional memory competition, we also employed pattern similarity of encoding-retrieval overlap to investigate whether retrieval of an extinction memory reactivates patterns of neural activity associated with the formation of an extinction memory.

**Fig. 1.**
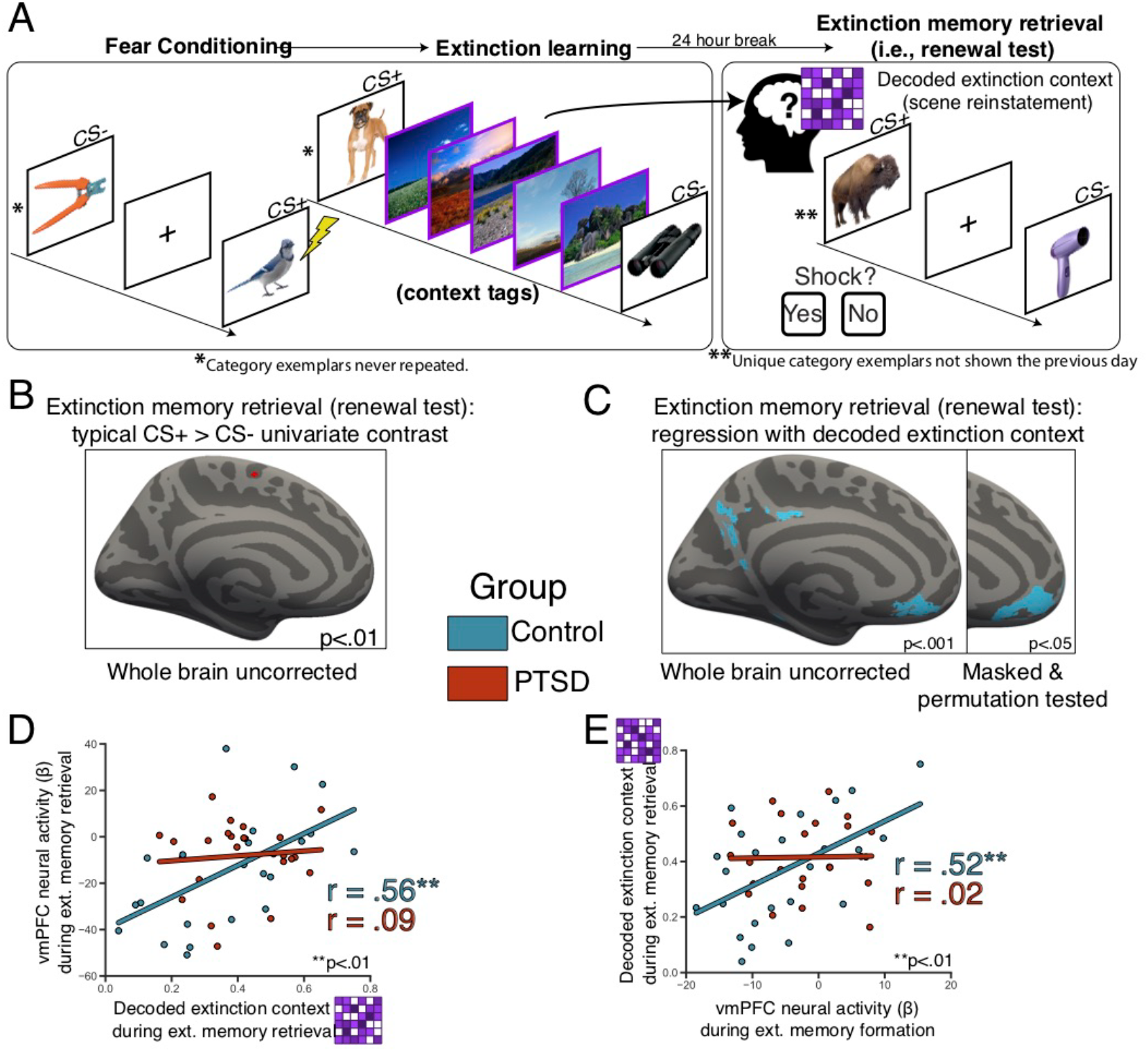
Extinction mental context decoding. **a.** Experimental Design. During fear conditioning 50% of CS+ co-terminated with a mild electric shock (US). During extinction, no shocks were delivered, and the ITIs were replaced with natural scene context tags. 24 hours later, participants were placed back into the scanner and shown novel CS+/− images. MVPA classifier evidence on CS+ trials during the renewal test provided evidence for reinstatement of the mental context associated with scene images from extinction memory formation. **b.** A GLM contrast was fit modeling early (first 4 trials) of CS+ activity greater than CS-activity. Activation map is weakly threshold at uncorrected p>.01 in order illustrate the inability of this technique to capture involvement of the vmPFC in extinction retrieval processes. **c.** Left. Whole brain univariate-multivariate regression reveals significant relationship in controls only, display threshold at p<0.001 uncorrected. Right. vmPFC ROI analysis, group functional mask was applied, and data was permutation tested (N fold = 1000), displayed at threshold p<0.05. 68% of the masked voxels met permuted threshold. **d.** During the early renewal test, extracted univariate (CS+ > CS-) vmPFC activity correlated with decoded extinction context in controls (blue, r=0.56, p=0.004) but not PTSD (red, r=.09, p=0.68). **e.** Univariate activity in the vmPFC at the time of extinction memory formation on Day 1 also correlates with decoded extinction context on Day 2 in the PPA during renewal testing in the control group (blue, r=0.52, p=0.009), but not in PTSD (red, r=0.02, p=0.92).

## RESULTS

We developed an fMRI experiment designed to tag and track the encoding and retrieval of an extinction memory by identifying patterns of neural activity associated with the context in which extinction was learned (Fig. 1a). Day 1 consisted of two phases: category fear conditioning^13,14^ followed immediately by extinction with mental context tags inserted between trials. We adapted a category fear conditioning design, in which conditioned stimuli (CS) in each phase consisted of trial-unique (i.e., non-repeating) basic-level exemplars from one of two distinct semantic categories (pictures of animals and tools). During conditioning, images from one category (the CS+, animals or tools) were paired with an aversive unconditioned stimulus (US, electric shock to the fingers) at a rate of 50%. Images from the other category (the CS-, tools or animals, respectively) were never paired with shock. Participants then underwent extinction during which additional novel and unique CS+/− images were presented, all without shock.

Multivariate pattern analysis (MVPA) can reveal how mental context reinstatement organizes episodic memory retrieval^7–9^, in accordance with computational models of contextually mediated episodic memory retrieval^5,15^. To tag and track the extinction mental context, we injected natural scene images as “mental context tags” between each CS trial during extinction learning only. Scene-related activity should therefore be assimilated into a mental context representation specific to the formation of the extinction memory. The following day, participants were exposed to unique exemplars from the CS+/− categories in a test of extinction memory retrieval, i.e., a fear renewal test. Critical to the design of this experiment, none of the exemplars shown on Day 2 were seen the previous day during conditioning or extinction, and were therefore entirely unique basic level exemplars from the same semantic categories used during both fear conditioning and extinction the previous day. In this way, the stimuli themselves were threat ambiguous, as they shared a semantic association with CSs used during both fear conditioning and extinction the previous day. The basic level exemplars (e.g., a bird, a dog, a bison, etc.) used as CSs during each phase (fear conditioning, extinction, or the renewal test) were counterbalanced across subjects. Also important to this design, participants did not see any natural scene images during the fear renewal test on Day 2; thus, the protocol resembles an ABA renewal test. Because scenes were only presented at extinction the previous day, scene-related activity detected during the fear renewal test can be interpreted as reinstatement of the extinction mental context. Neural analyses focused on the early renewal test (first 4 CS+/− trials), consistent with extant fMRI studies of extinction retrieval in human neuroimaging^16,17^.

### Traditional univariate approach

To date, neuroimaging studies utilizing classical conditioning paradigms to investigate competition between emotional memories have largely focused on mass univariate analyses to make inferences about extinction memory retrieval. Specifically, these studies use GLMs contrasting CS+ related activity against CS-related activity. However, recent comprehensive fMRI meta-analyses of human fear extinction indicate the typical CS+ > CS-contrast is often insufficient to capture activity in the vmPFC, despite extensive neurobiological evidence from animal models showing this region is critical for extinction memory formation and extinction memory retrieval^18^. We likewise failed to detect vmPFC activity from the typical CS+ > CS-univariate contrast from fear renewal test in both healthy and PTSD participants, even at an extremely liberal statistical threshold (Fig. 1b). A failure to capture activity in the vmPFC using univariate approaches might incorrectly support an interpretation that the vmPFC does not participate in extinction memory retrieval in the human brain. The subsequent analyses overcome the limitations of the common univariate approach to studying extinction processes in humans to reveal the role of extinction mental context reactivation on vmPFC activity.

### Extinction mental context decoding

To track neural reinstatement of the extinction mental context, a machine learning classifier was trained to identify scene-related activity in the parahippocampal place area (PPA) using fMRI data from a separate perceptual localizer task^9^. This classifier was then used to estimate reinstatement of the extinction mental context on CS trials during the renewal test by quantifying neural scene-related evidence in the PPA. Results of extinction context reinstatement on CS trials during fear renewal showed a wide distribution of classifier evidence for neural scene reinstatement in both healthy and PTSD subjects. Notably, the variability in mean classifier evidence was similar across groups and not different between healthy adults and PTSD group (ind. two-tailed t-test, t=-0.94, p=0.35; **Supplementary Fig. 1**). We extracted classifier evidence from each subject in order to examine the relationship between the magnitude of mental context reinstatement and neural activity in canonical extinction neurocircuitry regions of interest during early renewal test (first 4 CS+/− trials; Fig. 1d), and we found that control subjects exhibited a positive relationship (r=0.56, p=0.004). A whole-brain regression analysis revealed a significant positive correlation between MVPA classifier evidence for scenes and univariate parameter estimates in the vmPFC during fear renewal (Fig. 1c). We also investigated whether the amount of neural scene evidence at renewal (Day 2) was related to vmPFC activity at the time of extinction memory *formation* (Day 1). There was a positive correlation in healthy controls (r=0.52, p=0.009) suggesting that the strength of extinction memory formation in the vmPFC is related to the magnitude of mental context reinstatement of that context in PPA at a later test of extinction memory retrieval in a different context (Fig. 1e). Importantly, vmPFC neural activity was not correlated between extinction learning and renewal test for controls (r=0.02, p=0.93), indicating that vmPFC neural activity from these timepoints is independently related to the degree of extinction mental context reinstatement on Day 2.

Unlike healthy controls, PTSD participants did not show a relationship between the magnitude of reinstatement of the extinction context in the PPA and extinction-related univariate activity in the vmPFC or hippocampus (Fig. 1c-e), despite similar levels of classifier evidence of context reinstatement in the PPA as healthy controls (**Supplementary Fig. 1**). Thus, extinction neurocircuitry in PTSD may be compromised such that extinction retrieval processes are insensitive to the magnitude of extinction context reinstatement. This is consistent with neuroimaging data that show dysfunction both the structure and function of the hippocampus and vmPFC in PTSD^10^.

Human neuroimaging studies have also failed to capture extinction related activity in the amygdala^18^, despite the importance of amygdala inhibition for successful extinction retrieval^19^. In the current study we similarly did not observe any significant univariate activity, or a relationship between extinction context reinstatement and amygdala activity (**Supplementary Figs. 2 & 3**). However, given the well-defined neurocircuitry of extinction retrieval we hypothesized that reinstated context might have an *indirect* effect on amygdala activity through upstream inputs. In order to evaluate possible indirect effects, we used a non-causal mediation analysis within a bootstrap framework in which neural activity in the amygdala was entered as the outcome, reinstated mental context as the predictor, and neural activity in vmPFC and hippocampus were both used as parallel mediators^20,21^ (Fig. 2). The correlations between the mediators and the predictor and outcome met necessary assumptions (**Supplementary Table 1 & Supplementary Fig. 2**). We found that, in controls only, reinstated context interacts with both the vmPFC and hippocampus resulting in significant intervening effects on amygdala activity (path a_1_b_1_=-26.7, p=0.02, a_2_b_2_=23.33, p=0.01). Even though PTSD subjects had slightly higher overall fMRI activation in the hippocampus and amygdala (**Supplementary Fig. 3**), there were no significant intervening effects (path a_1_b_1_=-1.06, p=0.79, a_2_b_2_=3.96, p=0.78). Thus, in this model, reinstated extinction context in the PPA is signaled in the vmPFC and hippocampus, and that information is relayed to the amygdala, consistent with current understanding of the directionality of the extinction circuit from rodent neurophysiology^11,19^. In sum, neural reinstatement of an extinction learning context appears to be related to neural activity in regions associated with context-dependent extinction retrieval.

**Fig. 2.**
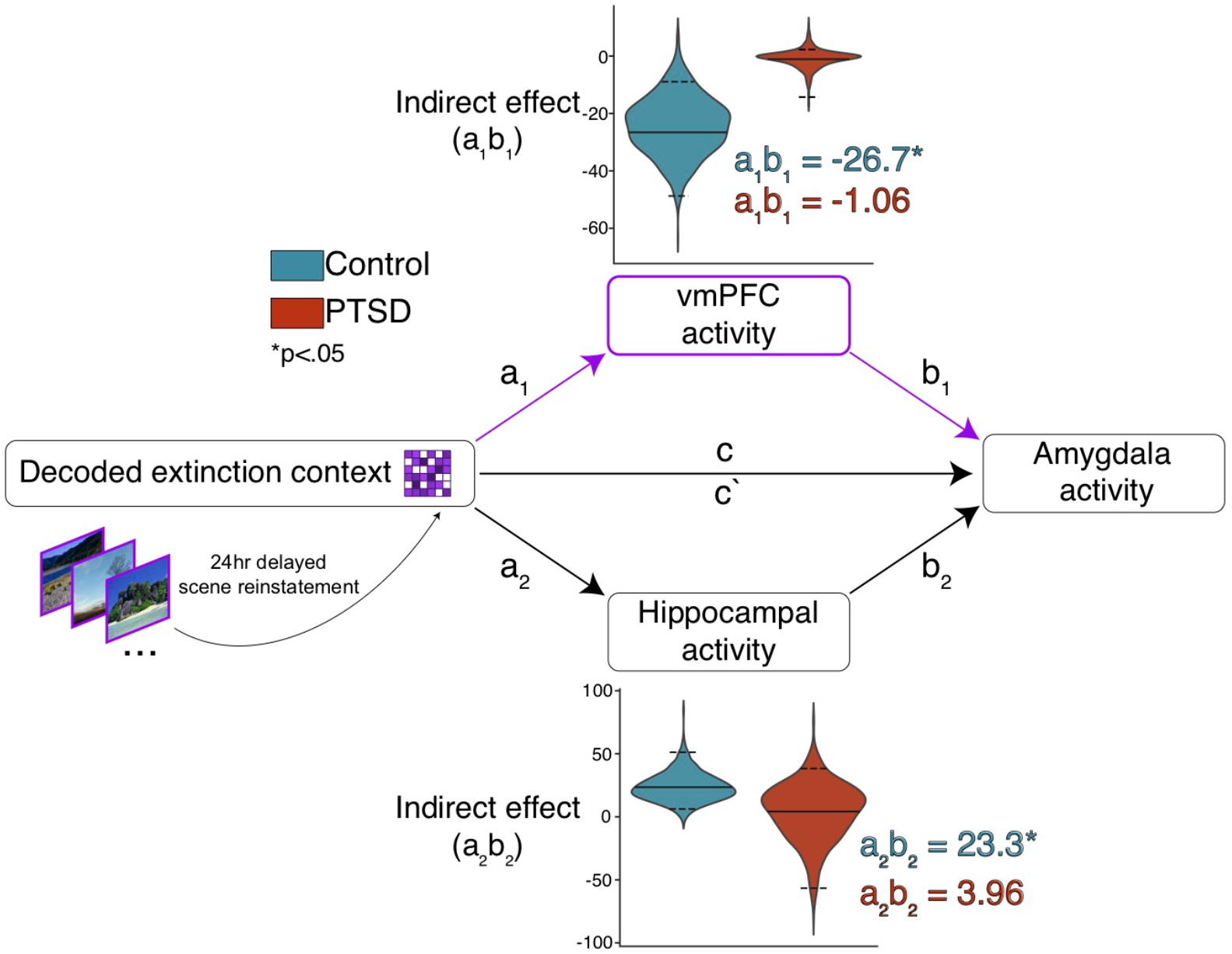
Reinstatement of extinction context indirectly effects amygdala activity through the vmPFC and hippocampus. Non-causal mediation analysis was performed separately for the control and PTSD groups. Full results for all paths are presented in **Supplementary Table 1.** No significant total effect (c) or direct (c`) effect was observed in either group. We tested if decoded extinction context exerts a significant indirect effect through either the vmPFC or hippocampus. Violin plots display distributions of indirect effects for vmPFC (a_1_b_1_) and hippocampus (a_2_b_2_) obtained from bootstrapping (N boots = 1000). Solid lines indicate mean indirect effect, dashed lines above and below indicate bounds of 95% confidence interval. The vmPFC indirect effect significantly differed from zero for controls (p=0.02) but not PTSD (p=0.78). The hippocampal indirect effect significantly differed from zero for controls (p=0.01) but not PTSD (p=0.79).

If reinstatement of the extinction mental context facilitates retrieval of the extinction memory, it might also help guide *behavior* when a novel and ambiguous CS is encountered. Accordingly, we assessed the relationship between explicit threat expectancy (**Supplementary Fig. 4**) and contextual reinstatement during the first CS+ trial of the renewal test, when the threat value of CS+ was the most ambiguous. We probed contextual reinstatement in the PPA based on subjects’ threat expectancy rating on the first CS+ trial (2 alternative forced-choice). The expectation that the CS+ would not be paired with a shock was taken to reflect the explicit retrieval of the extinction memory, as opposed to relapse of the fear memory. Healthy adults who perceived the first CS+ presentation in the renewal test as safe (N=15) had significantly higher neural reinstatement of the extinction context compared to those who anticipated threat (N=9). A Response by TR mixed ANOVA in controls reveals a main effect of response (F=4.12, p=0.05), and follow up t-tests show subjects who perceived the stimulus as safe had significantly higher scene evidence at TRs=- 2,0 relative to stimulus onset (TR=-2 two-tailed ind. t=2.16, p=0.04; TR=0 t=2.24, p=0.037; Fig. 3a). In contrast, PTSD participants showed no relationship between behavioral threat expectancy and extinction context reinstatement (Response by TR mixed ANOVA: main effect of TR only (F=6.57, p=0.0006), despite similar levels of neural scene reinstatement to controls. Thus, the magnitude of mental context reinstatement was associated with retrieval of an explicit extinction versus fear memory in healthy controls, but the magnitude of contextual reinstatement was unrelated to explicit extinction memory retrieval in PTSD.

**Fig. 3.**
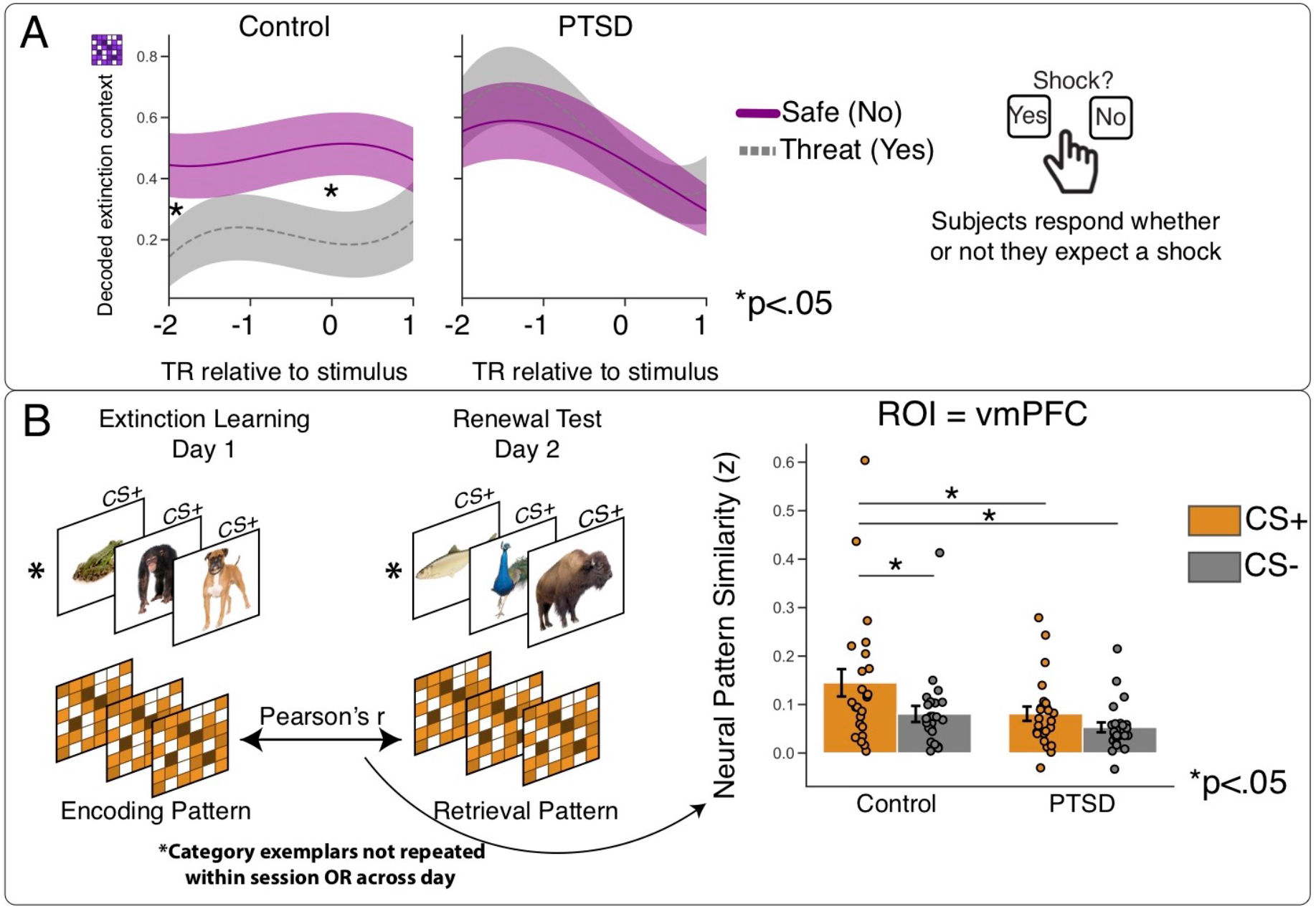
Decoded extinction mental context related to conscious threat expectancy. **A.** Subjects responded “Yes” or “No” if they expected a shock, and groups were split by first CS+ threat expectancy. Control subjects that perceived the stimulus as safe (did not expect a shock; N=15) had significantly higher classifier evidence for scenes in the PPA compared to subjects that perceived threat (expected a shock; N=9). No differences were observed for PTSD subjects who did (N=12) or did not perceive (N=12) a threat. Data are shifted 2 TRs to account for hemodynamic response and interpolated across timepoints for smoothness. Shaded regions indicate +/− 1 s.e.m. around mean (darker line). **B**. Extinction encoding-retrieval pattern similarity analysis. Left: CS+/− evoked patterns during extinction learning were correlated with never before seen novel CS+/− evoked patterns from the fear renewal test in the vmPFC. Right: Fischer-Z transformed CS+/− neural patterns for control and PTSD groups. Error bars represent +/− 1 s.e.m.

A complementary approach to assess successful memory retrieval is to quantify the overlap in neural activity between encoding and retrieval^22^. In human episodic memory research, for instance, the match between neural activity at encoding and retrieval predicts memory performance^23^. We adapted a measure of encoding-retrieval overlap to estimate the fidelity of extinction memory retrieval in the vmPFC between healthy adults and PTSD participants. We focused on representational overlap of pattern similarity in the vmPFC based on both the importance placed on this region for encoding and retrieving extinction memories and our own evidence linking extinction mental context reinstatement and neural activity in this region. A pattern similarity analysis^24^ was used to probe the similarity of CS evoked neural patterns of activity in the vmPFC between extinction learning and retrieval. In order to evaluate extinction-learning specific patterns, we also calculated pattern similarity for trials from the CS-category across extinction learning and renewal test (i.e., encoding-to-retrieval) as a comparison (Fig. 3b). Note, unlike a traditional episodic memory analysis focused on encoding-retrieval activity to the same item, here patterns of activity were compared across days between perceptually unique exemplars from the same semantic category (i.e., animals and tools). Further, subjects here were not engaged in explicit memory retrieval; rather, fear renewal tests constitute a more subtle test of emotional memory retrieval between competing associative memories (fear or safety). Thus, successful extinction memory retrieval can be construed within the framework of cortical reinstatement or transfer appropriate processing by quantifying neural similarity between extinction learning and extinction retrieval. The use of unique items across days also mitigates the potential role of mere perceptual overlap between stimuli used at encoding and retrieval. A CS type by Group mixed ANOVA revealed significant main effects of group (F=4.2, p=0.045) and condition (F=10.39, p=0.002). Planned t-tests revealed Control subjects displayed significantly greater encoding-retrieval similarity of CS+ extinction memories in the vmPFC compared to PTSD participants (two-tailed ind. t=2.01, p=.05, without outliers t=2.84, p=0.007). Healthy controls also showed a selectively enhanced similarity of evoked patterns of neural activity in the vmPFC for CS+ images over CS-(two-tailed paired t=2.74, p=.01), whereas PTSD participants show no such selectivity in increased pattern similarity between CSs (two-tailed paired t=1.71, p=.09). These data provide new evidence that patterns of neural activity in the vmPFC during the formation of extinction memories are later reinstated at extinction memory retrieval in healthy adults to threat ambiguous but conceptually related stimuli. In addition, control subjects also show higher overall neural pattern similarity in the hippocampus (CS by Group mixed ANOVA main effect of group; F=4.12, p=0.04; **Supplementary Fig. 6**), and selectively higher similarity for CS+ patterns in the PPA (two-tailed ind. t=2.12, p=0.04, **Supplementary Fig. 6**) as compared to PTSD participants. Overall, healthy adults appear to reinstate neural patterns associated with extinction learning as compared to PTSD participants, perhaps promoting successful retrieval of extinction memories outside the extinction context.

## DISCUSSION

This study presents new evidence that reinstated episodic mental context helps resolve the competition between competing emotional memories of learned threat and safety. Understanding the context dependent nature of emotional memory retrieval competition helps advance our understanding of affective disorders, for which dysregulated contextual processing may be a core feature. In healthy adults, the mnemonic signature of extinction context reinstatement positively correlated with activity in the vmPFC and hippocampus. Neurobiological research shows these regions are critical for encoding and retrieving extinction memories, but evidence from human fMRI studies utilizing typical univariate approaches only weakly support translation of these findings from rodents to humans. Here, activity in these regions was only revealed by accounting for the magnitude of extinction mental context reinstatement, as the standard CS+/− contrast did not reveal activity during the renewal test in either controls or PTSD (Fig. 1b). Despite a well-recognized role for the vmPFC in extinction processes^11^, failures to observe vmPFC activity in human fear extinction are common^18^. By utilizing multivariate fMRI analyses to explicitly account for reinstated episodic context during competitive emotional memory retrieval, we show extinction-related processing that is otherwise undetectable using standard neuroimaging approaches. Additionally, our non-causal mediation analysis showed that episodic mental context reinstatement has an indirect effect on amygdala activity through the vmPFC and hippocampus. The role of these structures in the retrieval of emotional memory in both animal models and humans is well known^11^. By incorporating multivariate information from our experimentally injected context tag, we provide new evidence that memory retrieval between competing emotional experiences is resolved by the amount of episodic context reinstatement.

Our study utilizes a category conditioning paradigm which allows for inferences about both episodic and associative memory^25^. On each trial subjects are presented with a trial-unique (i.e., non-repeating) basic level exemplar from one of two semantic categories. Subjects thus engage in episodic encoding of each unique category exemplar, and over time the semantic category acquires emotional associative meaning. This paradigm is well suited for the integration of mental context tagging. Mental context theories posit that encountering novel stimuli drives the formation of an episodic context neural representation^26^. Here, during the subsequent fear renewal test, subjects are presented with entirely new, and therefore threat ambiguous, stimuli and must evaluate the higher order semantic properties of each stimulus in order to determine which mental context, and related emotional association, to retrieve. By experimentally imbuing the extinction context with natural scene images, we were able to determine that healthy subjects who reinstated the extinction mental context when encountering novel stimuli were more likely to activate canonical extinction neurocircuitry (Fig. 3a). Specifically, we provide evidence that an episodic context signal obtained from the PPA also supports the resolution of associative ambiguity in the vmPFC, hippocampus, and amygdala (Fig. 2). This result contributes to the idea that associative memory competition is resolved in part by past and present episodic situational details^1^.

It is worth considering that the nature of the context tag we used here (scene images) were used because scene images reliably activate the PPA more strongly than other kinds of visual stimuli. Another category that reliably activates regions of sensory cortex (e.g., pictures of faces, odors, tactile stimuli) could presumably be used to equal effect. The ability to tag and track reinstatement of a multisensory naturalistic context could advance the utility of this protocol. For example, treatments for affective disorders and addiction based on associative learning theory contend that successful treatment establishes new associative memories that outcompete and inhibit retrieval and expression of maladaptive memories. Decoding reinstatement of an environment associated with formation of new adaptive associations might help measure the efficacy of a clinical treatment designed to prevent relapse of fear or addiction. A further advancement of these findings would be to enhance the neural representation of a context of safety using closed-loop neurofeedback with the goal of biasing memory retrieval toward safety and away from fear.

Pattern similarity analysis ^24^ provided a unique tool for assaying the strength of the extinction memory representation in the vmPFC. The overlap in neural activity between encoding and retrieval is widely thought to modulate the strength of episodic memory. We found that patterns of neural activity at fear renewal matched patterns from extinction learning, which might reflect a possible mechanism by which healthy adults retrieve the memory trace of extinction.^27^. Importantly, the items presented during fear renewal were threat ambiguous, and did not perceptually match the items encoded during extinction the previous day (i.e., the items in both phases were from the same category, but were unique exemplars). It is therefore noteworthy that encoding-retrieval overlap was greater for CS+ items as compared to the control CS-items, thereby indicating a unique method for measuring the strength of extinction memory retrieval by quantifying the overlap in pattern similarity between extinction learning and extinction retrieval.

As expected, PTSD participants displayed different pattern of results than healthy adults. That is, the PTSD symptomatic group did not show any discernable relationship between reinstated mental context and vmPFC activity during competitive emotional memory retrieval. Additionally, the mediation analysis showed that PTSD participants’ degree of episodic contextual reinstatement had no effects on amygdalar function via the vmPFC during competitive emotional memory retrieval. These results confirm earlier research highlighting dysregulated vmPFC activity in subjects with PTSD during the processing of contextual information^28^. In this symptomatic group reinstated episodic context had no relationship to conscious behavior, and PTSD participants displayed significantly lower extinction learning-retrieval neural pattern similarity as compared to healthy controls. These results suggest that PTSD participants were not utilizing the same memory retrieval processes during a period of threat ambiguity, which further supports the idea that contextual processing deficits are a pathogenic marker at the core of PTSD^4^.

Taken together, this study provides new evidence that multivariate neural signatures of contextual reactivation and representational overlap of extinction memories during encoding and retrieval can reveal the context-dependent nature of emotional memory competition in humans. Reinstatement of encoding patterns associated with the spatiotemporal context in which extinction memories are formed may help balance retrieval of safety memories against the renewal of fear. These findings also highlight new approaches to investigate the context-dependent nature of fear extinction memory in the human brain, consequently helping bridge the substantial translational divide between fine-scale molecular imaging of activity-dependent neural tagging in animal neuroscience and human neuroimaging. That the link between neural reinstatement and extinction memory retrieval is compromised in PTSD suggests a potential target for a disorder characterized by dysregulated contextual processing and extinction retrieval deficits.

## Acknowledgments

We thank Michael Drew, Greg Fonzo, and Suzannah Creech for helpful discussions and comments.

## Author contributions

A.C.H, J.A.L.-P., and J.E.D. conceived of and designed the experiment; M.M. recruited fMRI participants; A.C.H and M.M. performed the fMRI experiments; A.C.H. analyzed data; A.C.H, J.A.L.-P., and J.E.D. wrote the manuscript.

## Online Methods

Further information and requests for resources should be directed to and will be fulfilled by the Lead Contact, Joseph E. Dunsmoor.

### Participants

48 members of the University of Texas and the wider community of Austin participated in both days of the experiment. Participants were compensated at the rate of $30 per hour. All participants provided written informed consent and procedures were in compliance with the regulations and standards of the Institutional Review Board of the University of Texas at Austin (IRB # 2017-02-0094). Half of the subjects (N=24) made up our control group, all of whom self-reported no history of psychiatric illness and no current treatment for any psychiatric disorder. Mean age 21 years (s.d. 2 years); 15 female, 9 male. The other half of the subjects (N=24) comprised our PTSD symptom group. This group responded to flyers seeking volunteers with Trauma or Anxiety. Subjects were initially screened by phone for self-reported current diagnosis of PTSD without a neurologic history or substance abuse history. Subjects who passed inclusion and exclusion criteria were then invited to appear for an in-person screening where they completed questionnaires assessing severity of PTSD (PTSD Checklist for DSM-5; PCL-5), Anxiety (Beck Anxiety Inventory; BAI), and Depression (Beck Depression Inventory; BDI) symptoms (**see Figures S7 & S8**). Each subject in the PTSD symptom group listed exposure to a traumatic event, i.e., Criterion A. Of the 24 subjects included in the analysis, 24 were self-reported to have PTSD; two self-reported obsessive-compulsive disorder as their primary disorder. Mean age of the PTSD symptom group was 26 years (s.d 4.7 years); 17 female, 7 male). Given the high co-morbidity for PTSD and substance abuse disorder, subjects in the PTSD symptom group were given a urine toxicology test just prior to going into the MRI. No subjects in the PTSD symptom group tested positive for illicit drugs.

### Experimental Design

The experiment consisted of two sessions with a 24-hour delay between sessions. Day 1 consisted of a baseline phase, fear conditioning, and extinction with a mental context tag (scene images presented between CS trials). Day 2 consisted of a renewal test, a recognition memory test for the CS exemplars from Day 1, and a perceptual MVPA localizer. Data from the baseline phase and the recognition memory test are not presented here. All phases of the experiment consisted of participants viewing images presented via E-Prime, while inside the scanner and making responses on a 4-button box with their right hand. CS+/− stimuli were images of animals and tools. Images were obtained from www.lifeonwhite.com or publicly available resources on the internet. Each CS was a different basic level exemplar with a unique name (i.e., there were not two different pictures of a chimpanzee). Threatening or typically phobic stimuli were excluded (e.g., spiders, snakes, knives). For conditioning, extinction, and renewal all CS+/− were displayed for 4, 4.5, or 5s and were followed by 5, 6, or 7s ITI. During these phases, participants responded on every trial whether or not they expected a shock via responding on a button box. Subjects were uninstructed about the CS-US contingencies, and were told that if they paid attention they might learn an association between the images and the shock. Skin conductance responses were collected via electrodes placed on participants’ left palm and measured with a BIOPAC MP100 System (Goleta, CA) (**see Figure S5**). Conditioning consisted of 24 stimulus presentations from both CS+/− categories. 12 CS+ presentations (50% reinforcement) co-terminated in mild electric shock delivered to the index and middle finger of the participants left hand. Shock intensity was calibrated for each participant to a level described as “highly annoying and unpleasant, but not painful”. CS+ category was counterbalanced across participants. Extinction consisted of 24 stimulus presentations from both CS+/− categories, and no shocks were delivered. During all ITIs of extinction, the fixation cross was replaced by a stream of natural scene images displayed for 1 s each with 5, 6 or 7 scenes per ITI. 24hrs later, participants were brought back to the scanner, both sets of electrodes were attached, and told that the experiment would continue as before. The renewal test consisted of 12 presentations of each CS+/− category, and no shocks were delivered. Behavioral and neural analyses of the renewal test were focused *a priori* on the first 4 CS+ and 4 CS-trials. Focusing on these early trials is consistent with other current human neuroimaging studies of extinction retrieval ^16,17^. This is because the early trials reflect retrieval and later trials might reflect re-extinction or additional extinction learning. Following the renewal test, participants completed 2 runs of a perceptual localizer. Participants were shown a stream of a unique category of images (1s on, 1s off) including animals, tools, natural scenes, indoor scenes, and phase-scrambled scenes. Images were shown in blocks of 8, and participants were told to respond if they detected any duplicate image in a block (1 duplicate / block). After exiting the scanner participants completed various mood surveys (**see Supplementary Figure 9**).

### Image acquisition

Scanning was completed using the Siemens Skyra 3T Human MRI scanner located at the Biomedical Imaging Center at the University of Texas at Austin. Functional data were acquired with a 32-channel head-coil. Functional image resolution was 3mm isotropic voxels (TR = 2000ms, TE= 29ms; FoV = 228mm; n(Slices) = 48) A multi-band factor of 2 was used with AC/PC auto alignment. An additional high resolution T1-weighted 3D MPRAGE scan (TR=1.9s, 1mm isotropic voxels) was collected to aid in registration and region-of-interest definition.

### Image preprocessing

Images were prepared using a combination of FSL (Oxford Centre for Functional MRI of the Brain (FMRIB) Software Library, ANTs (Advanced Normalization Tools), FreeSurfer, and in-house Python scripts^29–31^. DICOM conversion was accomplished using dcm2niix^32^. Skull stripping was accomplished via FreeSurfer recon-all. Functional runs were motion-corrected using FSL mcflirt, and then all functional runs were co-registered to the T1 structural image. In preparation for multi-variate pattern analysis, time series data had linear trend removed, were high-pass filtered (sigma=128s), and z-scored. Representational similarity and trial unique neural activity analyses required trial estimate LS-S style beta images^33^, which were estimated using FSL and in-house python scripts. When provided, GLM parameter estimates were computed using FSL FEAT fMRI analysis of the motion corrected data and included prewhitening, high-pass filters, and registration to MNI152 space via FLIRT with 12 degrees of freedom.

### Statistical Analysis

A combination of parametric and non-parametric resampling analyses were used to statistically evaluate all of the above results. Where mentioned, outliers were defined by data that are beyond [1.5*Inter Quartile Range] in either direction. The type and value of all calculated test statistics and significance values are reported in the text and figure captions.

### ROI selection

Extinction network ROIs (hippocampus, amygdala, vmPFC) were chosen *a priori* based on their known role in extinction and renewal. At the subject level ROIs were defined anatomically using the relevant FreesSurfer parcellations (“mOFC” was used for vmPFC). At the group level, the vmPFC was defined functionally via GLM parameter estimates of CS- > CS+ activity during conditioning in a model that contained a regressor for the US. A cluster of voxels was selected manually after thresholding the whole brain image at p=.001, uncorrected. For MVPA decoding of the mental context tag, a group parahippocampal place area ROI was also functionally defined following the procedure of Bornstein et al., 2017. GLM parameter estimates of [Scenes > Scrambeled Scenes | Objects] were obtained for each subject and threshold at p=.001, uncorrected. These subject maps were then binarized and stacked. A cluster corresponding to the parahippocampal place area (PPA) was selected based on the criteria that it show activation in 80% of subjects. The group PPA mask was registered to each subject from MNI152 space using FSL FLIRT with 12 degrees of freedom.

### Multi-variate pattern analysis

MVPA decoding was accomplished using the Sklearn Logistic Regression classifier in Python ^34^. The classifier was trained to detect natural scene images vs. scrambled scene images in the PPA. Classifier sensitivity was assessed via cross validation of the two localizer runs (mean classifier ROC AUC = 0.91, s.e.m. = 0.03). To test for extinction mental context reinstatement, the classifier was trained on scenes vs. scrambled scenes during the localizer, and then used to obtain classifier evidence (probability estimates) for scene-related activity during the renewal test. Classifier evidence for scenes was used to operationalize extinction mental context reinstatement. Pattern similarity analysis ^24^ was achieved using in-house Python scripts. Trial beta estimates from the early renewal test and extinction learning were weighted by overall parameter estimates from the relative timepoints to reduce noise, and a Pearson correlation was obtained between the two resulting sets of images. In order to test for the reliability of these correlations at the group level, individual subject r values were Fischer-Z transformed before statistical testing.

### Data and Code Availability

All deidentified behavioral and neuroimaging data, as well as all custom python analysis code are available upon request to J.E. Dunsmoor.

