## Supplementary Figures and Tables for "Neural reinstatement guides context-dependent emotional memory retrieval"

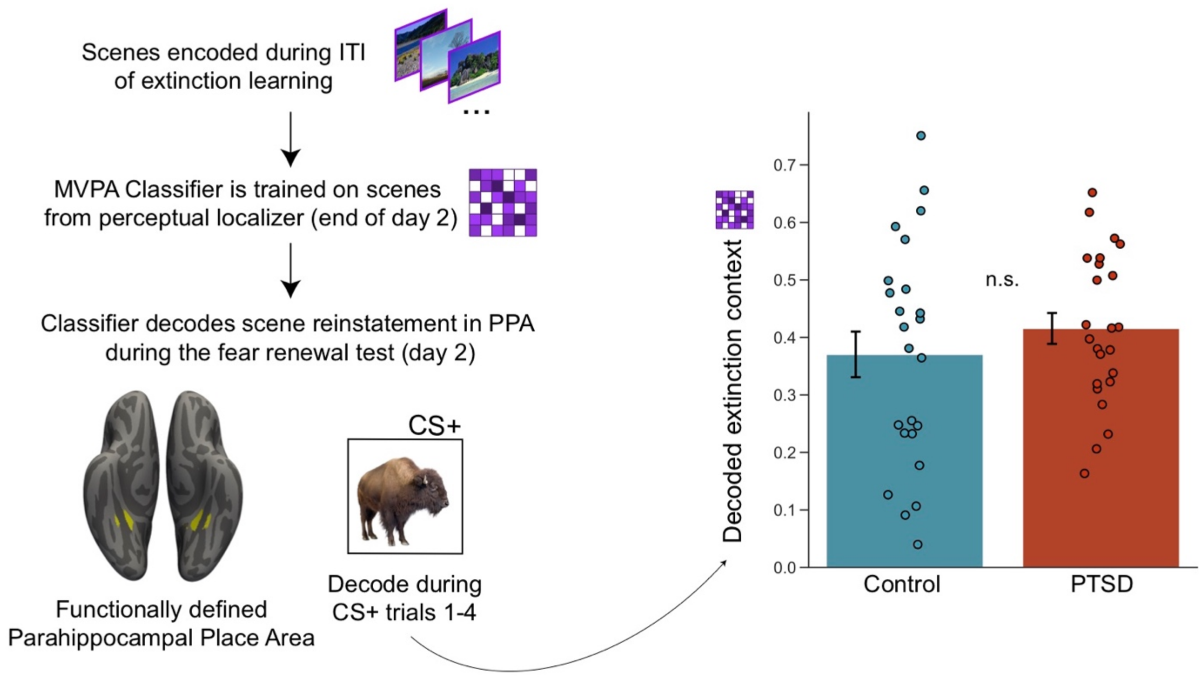


**Supplementary Fig. 1. Overview of extinction mental context decoding.** Participants encode task-irrelevant scene images during the inter-trial-intervals (ITIs) of extinction learning at the rate of 1 scene / second. Scene images were selected to be void of man-made structures and live subjects. A linear classifier (sklearn logistic regression) was trained on scene images and phase scrambled scene images collected during a perceptual n-back localizer at the end of day 2. The classifier was used to obtain estimates of scene related activity in the PPA during the first 4 CS+ trials of the day 2 renewal test. Total scene evidence did not differ between control and PTSD groups ($t=-0.94$, $p=0.35$). Error bars indicate +/- 1 s.e.m.

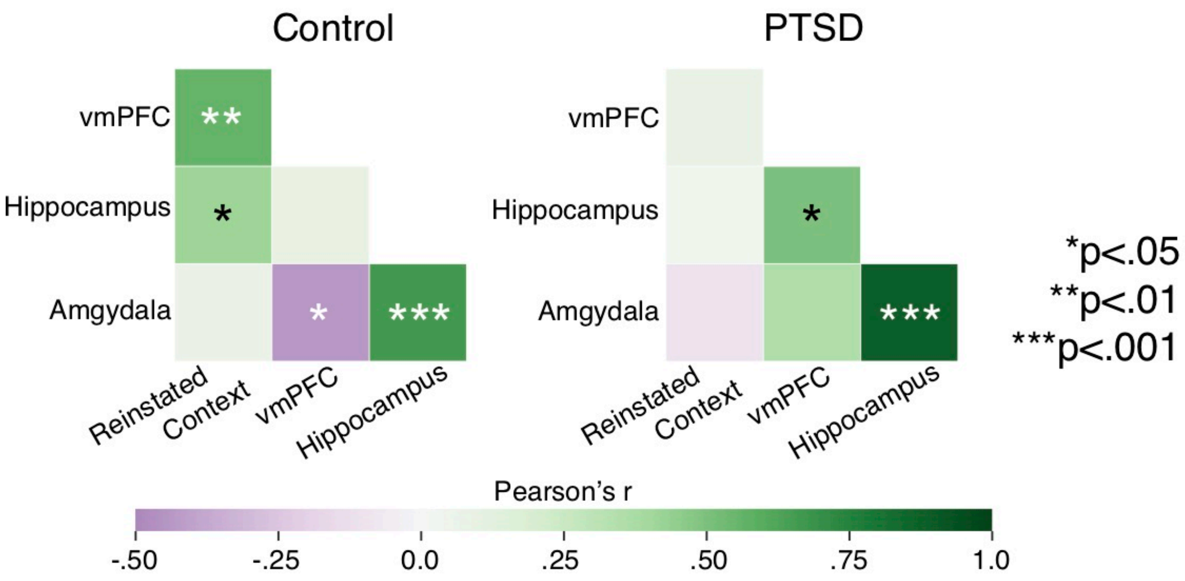


**Supplementary Fig. 2. Relationship between neural activity and reinstated extinction context in the extinction circuit.** Correlations between neural activity (univariate contrast of CS+ minus CS-) during the early renewal test and reinstated extinction mental context.

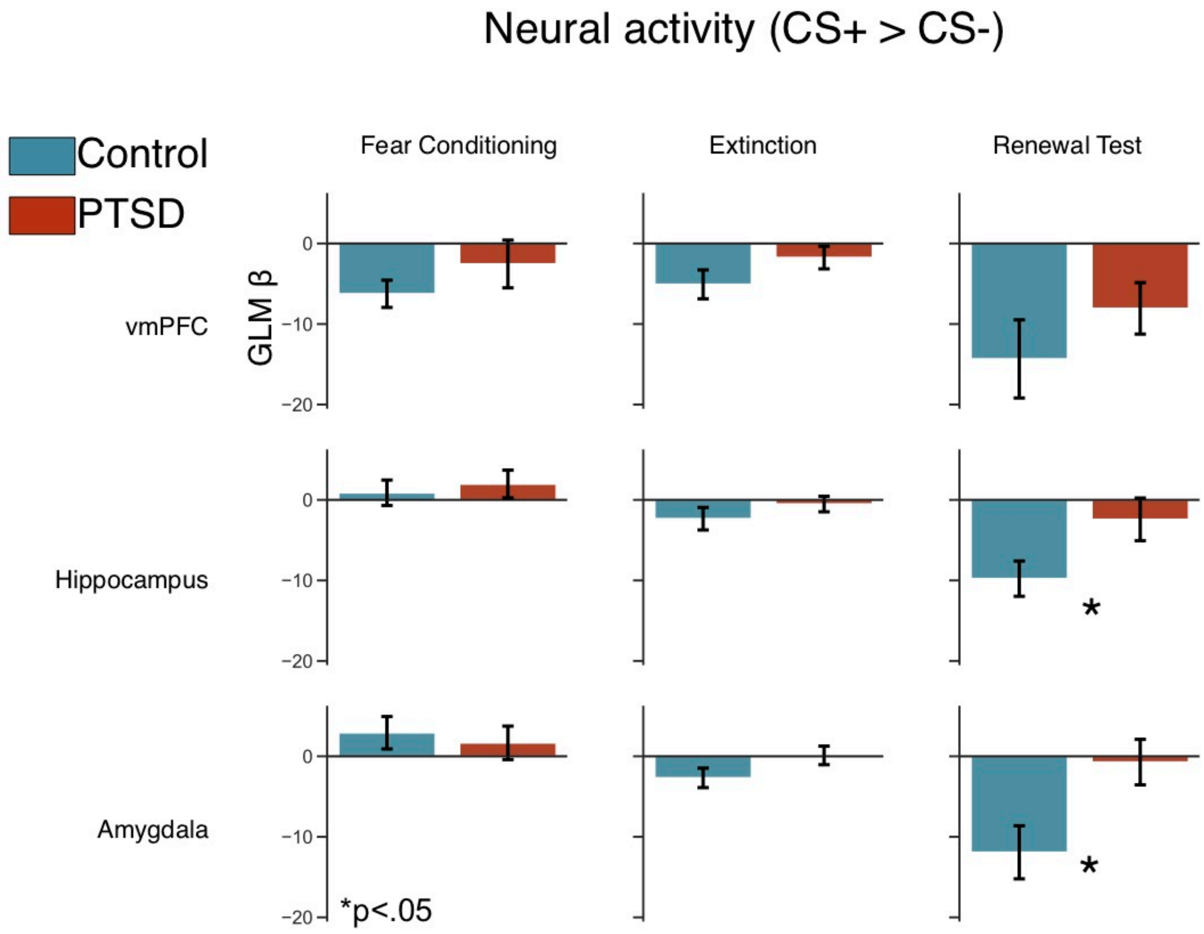


**Supplementary Fig. 3. Univariate results of CS+ minus CS- neural activity in extinction circuit across experimental phases.** Beta parameter estimates of the CS+ minus CS- contrast. Parameter estimates were extracted for each subject using anatomical Freesurfer labels registered into functional space (labels = hippocampus, amygdala, medial orbitofrontal cortex)**.** During the early renewal test, control subjects exhibited relatively diminished CS+ versus CS- activity in the hippocampus (ind. t=-2.14, p=0.037) and the amygdala (ind. t=-2.58, p=0.013). Error bars indicate +/- 1 s.e.m.

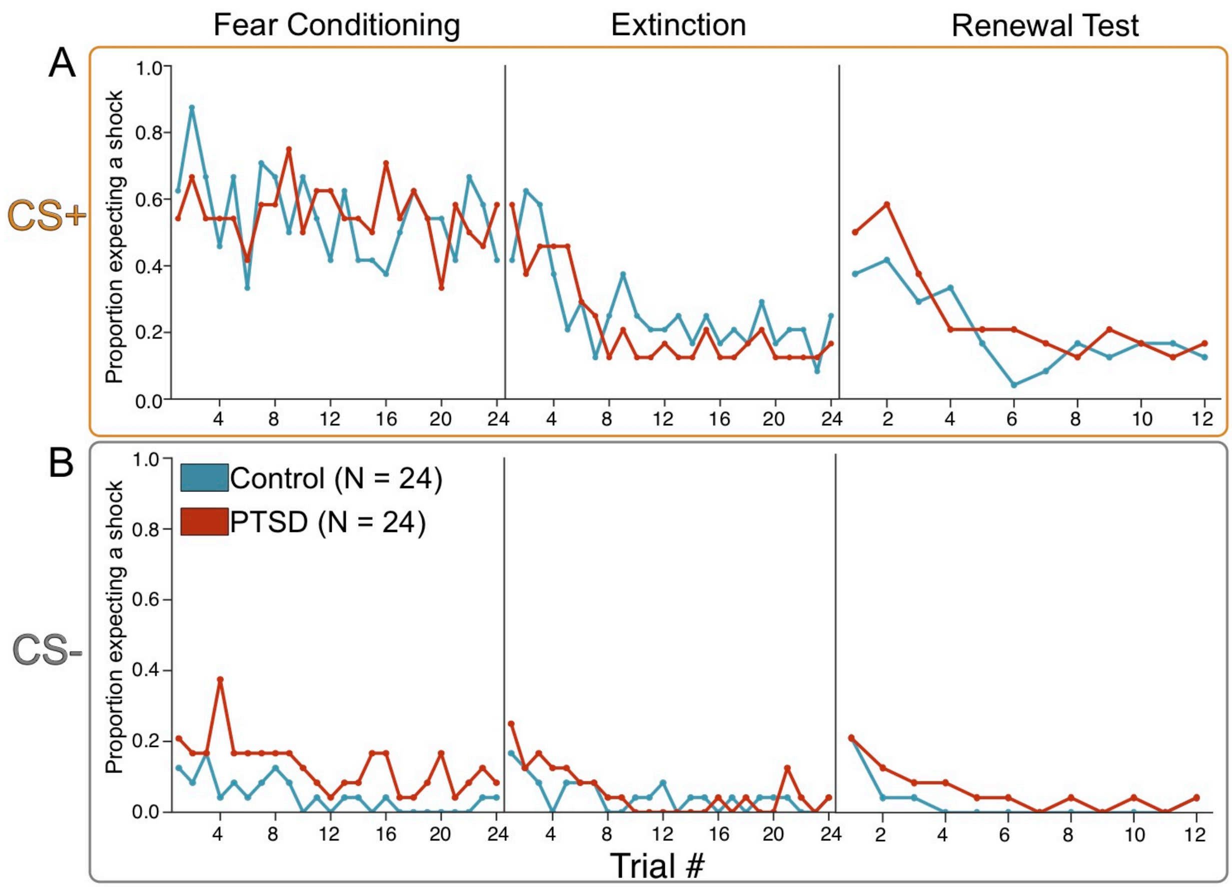


**Supplementary Fig. 4. Trial-by-trial shock expectancy behavior.** The proportion of subjects who responded that they expected a shock on each CS+ and CS- trial across phases for healthy controls and PTSD. Data reveal overall successful acquisition, extinction, and renewal of shock expectancy to the CS+ in both groups. The first trial of fear conditioning was a shock reinforced CS+ image, likely explaining why expectancy was initially low for the majority of subjects on the first CS- trial of fear conditioning.

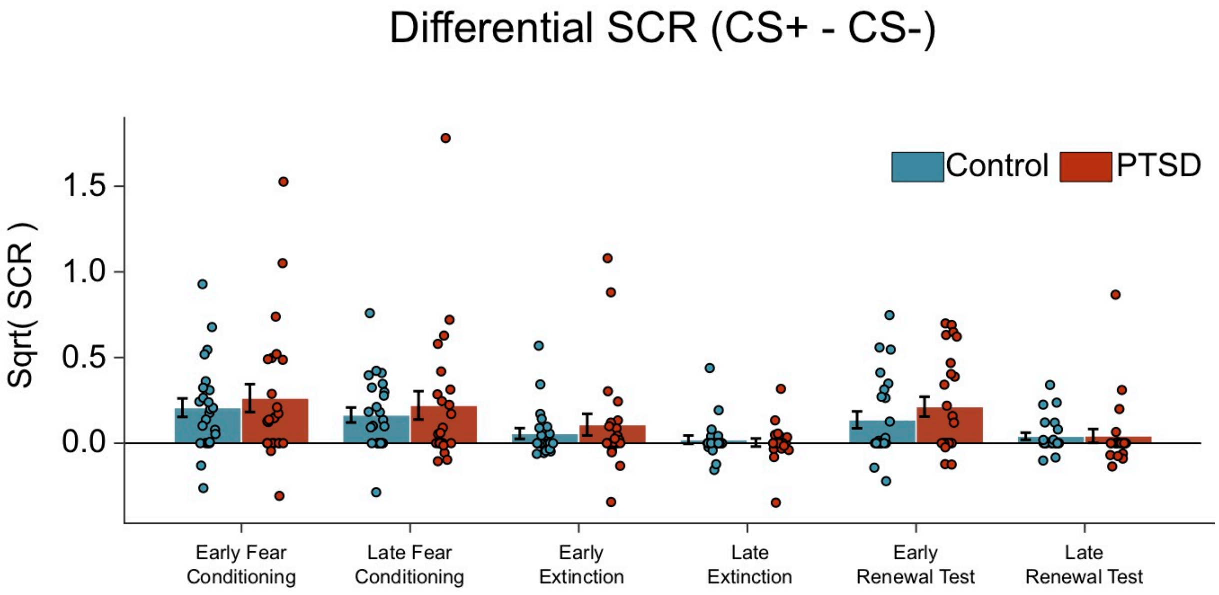


**Supplementary Fig. 5. Skin conductance response by experimental phase.** Differential (CS+ minus CS-) square-root normalized SCRs (CS+ - CS-). Early and late conditioning/extinction correspond to 24 total CS+/- trials each. Early renewal consists of 8 total CS+/-, late renewal 16 total CS+/-. Repeated measures ANOVA revealed a significant main effect of phase ($F=7.53$, $p=2e-5$). No effect of group was observed ($F=0.90$, $p=0.35$). These data show intact acquisition, extinction, and renewal of conditioned physiological arousal at the group level for healthy controls and PTSD. Error bars represent +/- 1 s.e.m.

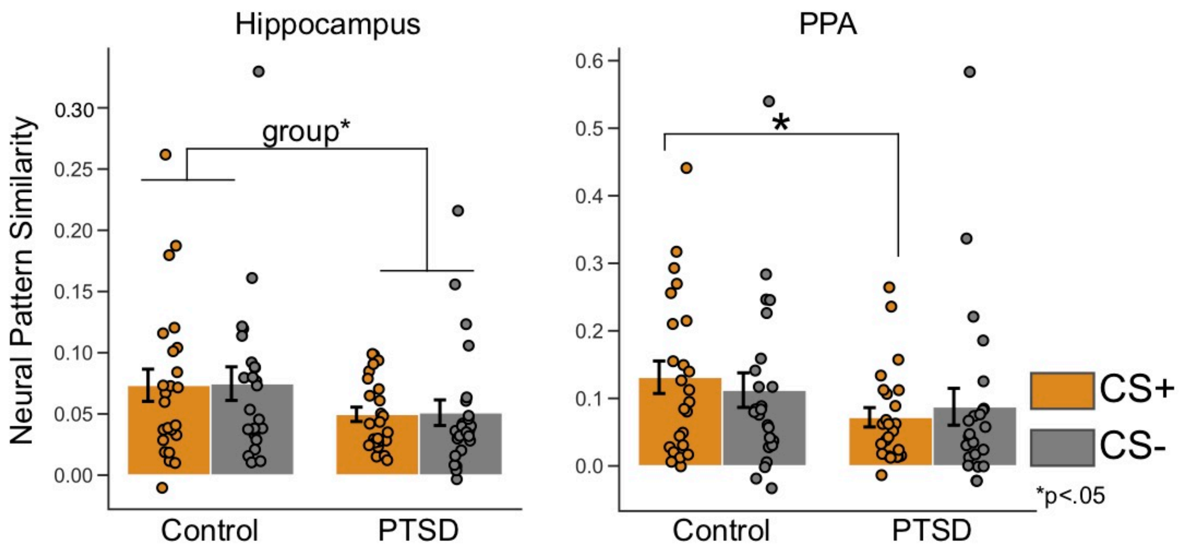


**Supplementary Fig. 6. Representational similarity analysis.** Procedure as described in text and **Fig. 3.** In the hippocampus, a CS type by Group mixed ANOVA revealed a main effect of group (F=4.12, p=0.048). In the PPA a two way mixed ANOVA relvelaed no main effects or interactions, but planned follow up t-tests show control CS+ pattern similarity is significantly higher than PTSD CS+ pattern similarity (ind. t=2.12, p=0.040). Error bars indicate +/- 1 s.e.m.

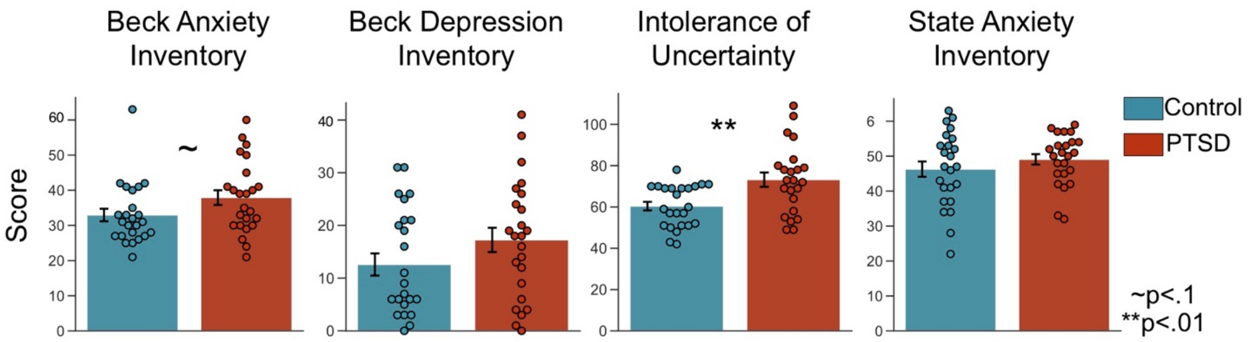


**Supplementary Fig. 7. Mood Surveys**. Compared to healthy control subjects, PTSD subjects scored significantly higher on the Beck Anxiety Inventory (t=1.81, p=0.077; without outliers t=-2.57, p=0.014), and the Intolerance of Uncertainty survey (t=3.16, p=0.003). There was no difference on state anxiety, which was assayed at the end of the experiment on day 1. Error bars indicate +/- 1 s.e.m.

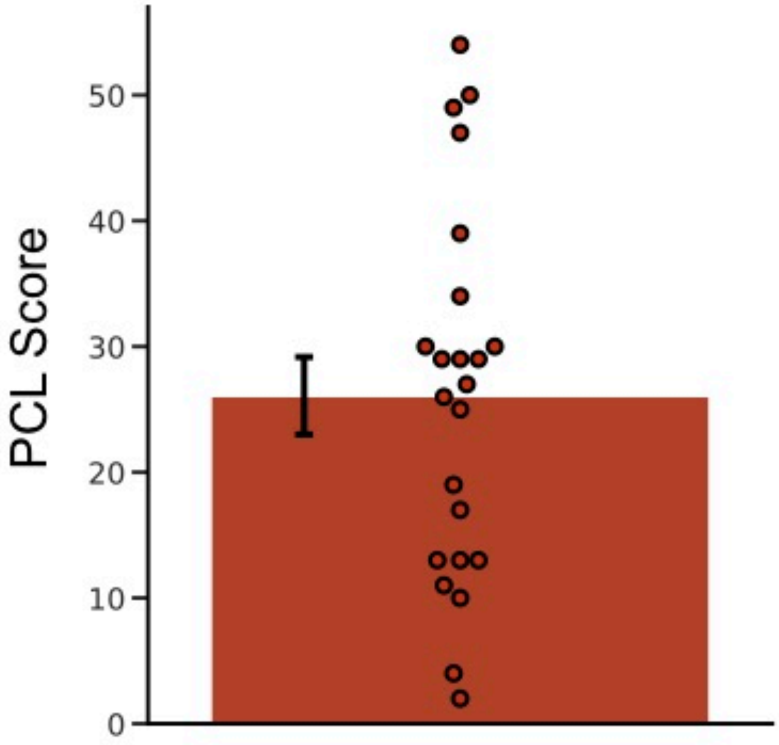


**Supplementary Fig. 8. Patient PTSD Checklist (PCL) scores.** Average PCL score for the patient group was 26.09. Error bar indicates +/- s.e.m.

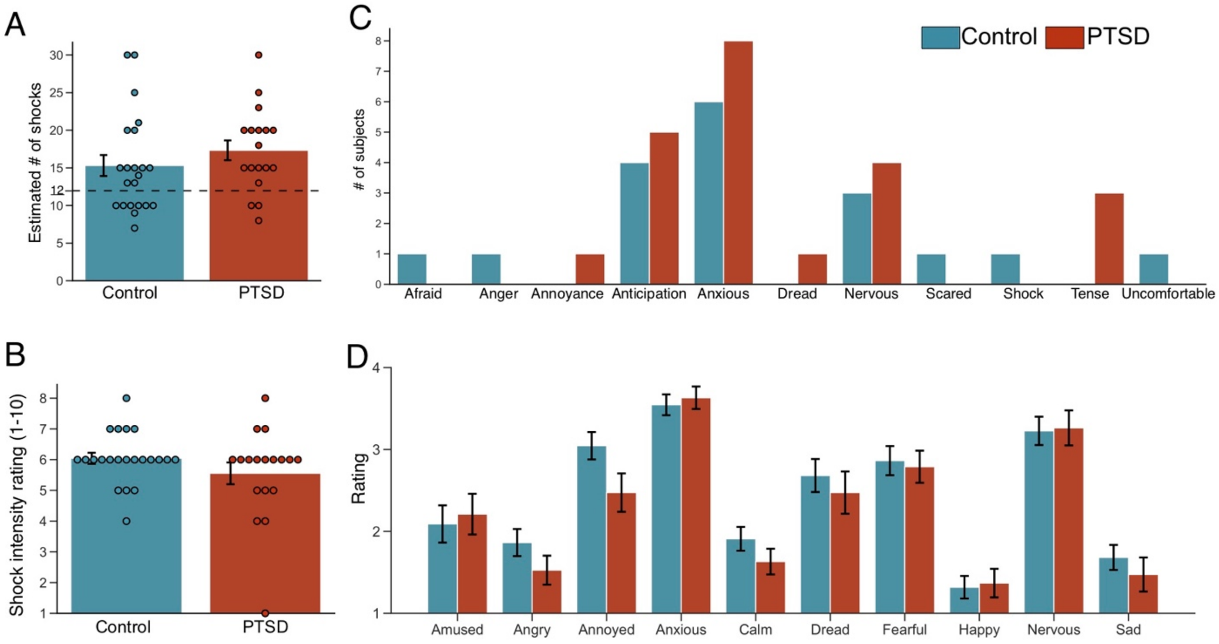


**Supplementary Fig. 9. Shock estimations and fear conditioning construct validity survey data.** At the end of day 1, subjects were asked to estimate the number of shocks they had received during the experiment (**A,** true # =12), and to retrospectively rate the intensity of the shock throughout the experiment (**B**). There were no differences in shock estimation or shock intensity ratings between healthy controls and PTSD. **C.** At the conclusion of the experiment, subjects were asked to respond with a single word that best described how they felt during the experiment. The purpose of this question was to address the construct validity of fear conditioning as inducing an emotional state akin to fear, or a synonymous construct. Healthy controls and PTSD similarly responded with words like "anxious," "nervous," or closely related negative emotional words. **D** Subjects then rated a series of words on how well each word described the emotion they felt when they expected a shock. (1 = not at all reflective to 4 = highly reflective). Again, healthy controls and PTSD similarly endorsed negative emotion words related to feeling anxious, nervous, or fearful. Error bars indicate +/- 1 s.e.m.

| Control Group | | | | | |
| --- | --- | --- | --- | --- | --- |
| Path | Estimate | Standard Error | Bootstrapped confidence interval 2.5% | 97.5% | p-value |
| $a_1$ | 68.53** | 21.54 | 23.86 | 113.21 | 0.004 |
| $b_1$ | -0.35*** | 0.08 | -0.52 | -0.18 | 0.0003 |
| $a_2$ | 22.86* | 10.70 | 0.66 | 45.06 | 0.044 |
| $b_2$ | 1.08*** | 0.18 | 0.71 | 1.46 | $1xe^{-5}$ |
| $a_1b_1$ | -26.71* | 10.16 | -48.81 | -8.95 | 0.016 |
| $a_2b_2$ | 23.33* | 11.44 | 6.17 | 51.25 | 0.010 |
| c | 5.86 | 17.64 | -30.73 | 42.45 | 0.743 |
| c` | 9.24 | 13.38 | -18.67 | 37.14 | 0.498 |

| PTSD Group | | | | | |
| --- | --- | --- | --- | --- | --- |
| Path | Estimate | Standard Error | Bootstrapped confidence interval 2.5% | 97.5% | p-value |
| $a_1$ | 10.53 | 25.21 | -41.76 | 62.82 | 0.68 |
| $b_1$ | -0.11 | 0.10 | -0.31 | 0.09 | 0.27 |
| $a_2$ | 3.89 | 20.96 | -39.58 | 47.36 | 0.85 |
| $b_2$ | 1.02*** | 0.12 | 0.77 | 1.27 | $2xe^{-8}$ |
| $a_1b_1$ | -1.06 | 3.24 | -14.28 | 2.30 | 0.79 |
| $a_2b_2$ | 3.96 | 22.9 | -56.61 | 38.18 | 0.78 |
| c | -11.16 | 22.27 | -57.36 | 35.03 | 0.62 |
| c` | -14.07 | 9.85 | -34.61 | 6.47 | 0.17 |

**Supplementary Table 1. Summary of paths from non-causal mediation analysis.** Bootstrap mediation analysis was accomplished using the Python statistical analysis package pingouin.
